# Inaccurate fossil placement does not compromise tip-dated divergence times

**DOI:** 10.1101/2022.08.25.505200

**Authors:** Nicolas Mongiardino Koch, Russell J Garwood, Luke A Parry

**Affiliations:** Scripps Institution of Oceanography, University of California San Diego, CA, USA; Department of Earth and Environmental Sciences, University of Manchester, Manchester, UK; Earth Sciences Department, Natural History Museum, London, UK; Department of Earth Sciences, University of Oxford, Oxford, UK

**Keywords:** Fossils, paleontology, phylogenetics, time calibration, morphology, fossilized birth-death models

## Abstract

Time-scaled phylogenies underpin the interrogation of evolutionary processes across deep timescales, as well as attempts to link these to Earth’s history. By inferring the placement of fossils and using their ages as temporal constraints, tip dating under the fossilised-birth death (FBD) process provides a coherent prior on divergence times. At the same time, it also links topological and temporal accuracy, as incorrectly placed fossil terminals should misinform divergence times. This could pose serious issues for obtaining accurate node ages, yet the interaction between topological and temporal error has not been thoroughly explored. We simulate phylogenies and associated morphological datasets using methodologies that incorporate evolution under selection, and are benchmarked against empirical datasets. We find that datasets of moderate sizes (300 characters) and realistic levels of missing data generally succeed in inferring the correct placement of fossils on a constrained extant backbone topology, and that true node ages are usually contained within Bayesian posterior distributions. While increased fossil sampling improves the accuracy of inferred ages, topological and temporal errors do not seem to be linked: analyses in which fossils resolve less accurately do not exhibit elevated errors in node age estimates. At the same time, divergence times are systematically biased, a pattern that stems from a mismatch between the FBD prior and the shape of our simulated trees. While these results are encouraging, suggesting even fossils with uncertain affinities can provide useful temporal information, they also emphasise that paleontological information cannot overturn discrepancies between model priors and the true diversification history.

## Introduction

Establishing an accurate timeline for the diversification of life on Earth is integral to understanding major events in evolutionary history and linking these to the history of the planet. Phylogenies scaled to absolute time underpin our understanding of the interplay between biology and geology (Mills et al. 2022, Palazessi et al. 2022), the interactions between lineages across deep time (Strassert et al. 2021, Asar et al. 2022, Onstein et al. 2022), the movement of clades across the globe (O’Hara et al. 2019, Landis et al. 2022), and the tempo and mode by which ecological, molecular and morphological novelties have emerged (Jabłońska & Tawfik 2021, Coombs et al. 2022, Mongiardino Koch et al. 2022, Suissa & Friedman 2022). Two distinct sources of information are used to infer divergence times: character datasets for the species under study, which reflect their relative degrees of divergence; and temporal constraints that help translate these into absolute times (Donoghue & Benton 2007). Combined with various modelling assumptions regarding patterns of character evolution, rate variation, and lineage diversification (Warnock & Wright 2021), these data produce time-calibrated phylogenetic trees (i.e., chronograms or timetrees) whose node ages have been estimated using a molecular and/or morphological clock.

While various sources of temporal evidence can be used to calibrate a phylogenetic clock model, fossil occurrences are the most widely employed (Hipsley & Müller 2014). When applied to the nodes of a phylogenetic tree, fossil calibration points provide minimum bounds for divergence times, as the earliest occurrence of a lineage in the fossil record necessitates older origination times. The minimum ages of multiple nodes can be bracketed in such a way, providing local temporal information that can be combined with relaxed clock models that accommodate heterogeneities in evolutionary rates among branches (Kishino et al. 2001, Drummond et al. 2006). However, biological and geological processes can impair the degree to which fossil occurrences approximate the ages of cladogenetic events (Marshall 2019), as well as the overall accuracy of fossil-calibrated clocks (e.g., Budd & Mann 2020, Carruthers & Scotland 2020). While some of the limitations of node-based approaches can be overcome by using fossils to derive prior distributions for node ages (Yang & Rannala 2005), the (semi)arbitrary configuration of these calibration densities can also largely determine the results obtained (Ho & Phillips 2009, Warnock et al. 2011, Brown & Smith 2018, Moody et al. 2022). Furthermore, results can also be compromised by incorrect assumptions regarding the relationships between fossils and extant terminals (Lee 2009, Near et al. 2005, Parham et al. 2012), which are treated with absolute certainty in node dating analyses.

An alternative approach to extracting the temporal signal from the fossil record is to incorporate extinct taxa as terminals whose phylogenetic placement is inferred using morphological information (O’Reilly et al. 2015, Wright et al. 2022). While methods to calibrate trees using fossil tips have existed for several decades (Fisher 1992, Wagner 1998), modern Bayesian implementations (often referred to as ‘tip dating’) have become dominant in recent years. Early tip-dating efforts (Pyron 2011, Ronquist et al. 2012) made use of simplistic models of diversification (also known as tree priors), but subsequent studies have applied fossilised birth-death (FBD) processes (Heath et al. 2014) that explicitly model speciation, extinction and fossil recovery dynamics. The switch to these mechanistic models of diversification has generally resulted in improved estimates of divergence times (Matzke & Wright 2016, Arcila & Tyler 2017, May et al. 2021). Simulations have also revealed that tip dating outperforms undated Bayesian and parsimony approaches in terms of topological accuracy (Mongiardino Koch et al. 2021), while being relatively robust to several potential sources of error (Klopfstein et al. 2019, Luo et al. 2020, O’Reilly & Donoghue 2021).

Calibrating phylogenies using fossil terminals provides numerous benefits in addition to improvements in the accuracy of tree topology and divergence times. First, it does not require specified prior distributions on node ages, avoiding a number of subjective—yet often impactful—decisions. Furthermore, the implementation of FBD processes provides a means of encompassing living and extinct terminals within a unified model of diversification and sampling, allowing the recovery of ancestor-descendant relationships that are not accommodated by other methods (Gavryushkina et al. 2014), and permitting the straightforward integration of morphological, stratigraphic and molecular datasets for total-evidence dated inference (Zhang et al. 2016, Heath et al. 2017). The resulting topologies not only provide an adequate basis for macroevolutionary studies (Slater & Harmon 2013, Arcila & Tyler 2017, Mongiardino Koch 2021), but macroevolutionary dynamics can even be incorporated into the process of phylogenetic reconstruction (Simões et al. 2020, May et al. 2021, Wright et al. 2021). Finally, tip-dated inference makes a much more thorough and objective use of paleontological data by potentially allowing all fossil species to be incorporated, not just the oldest representatives of clades; and inferring, rather than enforcing, their phylogenetic positions (O’Reilly et al. 2015, Wright et al. 2022).

The simultaneous estimation of tree topology and divergence times in tip-dated analyses allows for a better integration of sources of uncertainty through the interaction of morphological and stratigraphic signals (Ronquist et al. 2012, King et al. 2017, King 2021). On the other hand, it also results in a direct link between topological and temporal accuracy that is absent from node-based methodologies. Given that fossil terminals constrain divergence times only in the path that connects them to the root of the tree, inaccurate fossil placements should misinform node ages in a manner that depends on the magnitude of topological error (Barido-Sottani et al. 2022). This is problematic, as morphological phylogenetics suffers from many shortcomings, including the subjectivity associated with character selection and scoring, the limited size of datasets, the overall difficulty in modelling the process and rate of morphological evolution, and the high prevalence of selective processes that overprint historical signals (O’Reilly et al. 2015, dos Reis et al. 2015, Donoghue & Zhang 2016, Simões et al. 2017, Keating et al. 2020, Mongiardino Koch & Parry 2020). Furthermore, fossils are affected by high levels of missing data, which can reduce the precision with which they can be placed (Nixon & Wheeler 1992, Wiens & Moen 2008), and may sometimes systematically bias their inferred phylogenetic position (Sansom et al. 2010, Sansom & Wills 2013). Several studies, both empirical (Turner et al. 2015, Spasojevic et al. 2021, Mongiardino Koch & Thompson 2021) and simulation-based (Luo et al. 2020, Mongiardino Koch et al. 2021), have revealed the difficulties associated with accurately placing terminals devoid of molecular data (such as fossils). Some have even suggested that this impacts the accuracy of inferred divergence times to such an extent that it potentially negates the other benefits of tip-dating (O’Reilly et al. 2015, Donoghue & Zhang 2016). To date, the nature and strength of the relationship between the accuracy of fossil placement and inferred divergence times in tip-dated analyses under morphological clocks has not been characterised. We do so here, using simulations to test the hypothesis that more accurate fossil placement improves estimates of divergence times.

## Methods

### Simulation approach

Our simulation approach follows the procedures described in Mongiardino Koch et al. (2021). Evolutionary histories, encompassing both morphological datasets and phylogenetic trees, were generated using the individual-based simulation framework TREvoSim v2.0.0 (Keating et al. 2020). This approach differs in a number of ways from other commonly-used simulation methodologies, namely: it does not employ any of the models routinely used for phylogenetic inference (i.e., Markov substitution models or birth-death tree models); it outputs paired character matrices and phylogenetic trees that are concurrently generated by the simulation process, rather than producing these separately; and it incorporates important biological concepts such as an estimate of fitness and a species concept (Garwood et al. 2019, Keating et al. 2020). This introduces a level of model misspecification that is likely pervasive in morphological phylogenetics under probabilistic methodologies (Wright & Hillis 2014), as well as better resembling empirical morphological datasets that have been modelled by adaptive evolution (Zou & Zhang 2016). Settings were identical to those used by Mongiardino Koch et al. (2021), as these were validated to replicate properties seen in twelve empirical morphological datasets that include fossil terminals, such as evolutionary rates, distributions of branch lengths, and levels of treeness (empirical datasets and a TREvoSim settings file can be found in the supplementary materials of Mongiardino Koch et al. 2021). Topologies incorporated a series of short internodes close to the root and followed by long unsplit branches giving rise to major extant clades. This pattern is reminiscent of ancient rapid radiations (Whitfield & Lockhart 2007), which pose numerous challenges for phylogenetic inference and dating. As previously, settings were chosen to produce datasets containing 500 binary characters and 999 terminals, of which an average of 150 were extant.

### Data manipulation

Subsequent processing steps were undertaken in the R programming environment (R Core Team 2021), using scripts provided in the Supplementary Material, and relying on packages *ape* (Paradis & Schliep 2019), *Claddis* (Lloyd 2016), *dplyr* (Wickham et al. 2021), and *phytools* (Revell 2012). Phylogenetic trees were checked for zero-length branches, and datasets with non-fully bifurcating topologies were discarded. Given that the chronograms simulated by TREvoSim are scaled to the number of iterations, we translated these to absolute time by assuming a 210.9 Ma old root, given the average timespan of the twelve reference empirical datasets, and rescaled branch lengths accordingly. Datasets were then subsampled by: a) eliminating fossils at random to produce a dataset size of 300 terminals, a process that mimics the incompleteness of the fossil record; and b) selecting 300 parsimony-informative characters at random, mimicking the process of character selection involved in matrix construction. Simulated datasets were further modified by imputing missing data. We counted the number of characters that were missing/inapplicable for each taxon in the empirical datasets employed, and combined these to derive realistic distributions of the proportion of missing data in both living and extinct terminals (Fig. S1). Values were then drawn from these distributions, and data was deleted at random from the simulated characters accordingly, producing datasets that varied widely in their levels of missing data (Fig. S2).

In order to evaluate the effect of increasing numbers of fossils on the accuracy of inferred divergence times, each dataset was subsampled one last time to 100 final terminals, varying the proportion of fossil and extant tips. Seven levels of fossil sampling were explored (5, 9, 16, 28, 44, 61, and 75 terminals), which were chosen as they represent approximately doubling numbers of fossil tips per internal node connecting extant taxa (0.05, 0.10, 0.19, 0.39, 0.80, 1.61, and 3.12, respectively). We generated 250 datasets for every level of fossil sampling, using each simulated dataset only once.

### Phylogenetic inference

All phylogenetic analyses were performed in MrBayes v3.2.7 (Ronquist et al. 2012b). Given the difficulties assessing the accuracy of divergence times when topology is being co-estimated, as nodes might only exist in a fraction of posterior trees, we used partial constraints to enforce a topological scaffold connecting all extant terminals. This relied on functions from *paleotree* (Bapst 2012), and enforced the true (i.e., simulated) relationships among extant terminals, while leaving the position of fossils unconstrained. This approach has been advocated by many to improve topological and/or temporal accuracy of tip-dated inference, and reflects the fact that the topological uncertainty of genomescale datasets is negligible compared to that of morphological matrices (Ronquist et al. 2012a, Slater 2013, Beck & Lee 2014, Crawford et al. 2015, Lee & Palci 2015, O’Reilly et al. 2015, Donoghue & Zhang 2016, O’Reilly & Donoghue 2016, Brown & Smith 2018, Mongiardino Koch & Thompson 2021, Darlim et al. 2022).

Uncertainty in tip ages was modelled using information from the Paleobiology Database (PBDB, https://paleobiodb.org/). The entire PBDB was downloaded and used to build a distribution of species longevities. Occurrences were grouped by species, and longevities defined as the time spanned between the minimum age of their first occurrence and the maximum age of their last one. Longevities longer than 100 Ma were filtered out, and the remaining data was found to approximate an exponential distribution with a rate parameter of 0.115 (estimated using ‘fitdistr’ in *MASS;* Venables & Ripley 2002), corresponding to a mean duration of 8.67 Ma (Fig. S3). Time intervals were generated by sampling from this distribution and assigning to fossils in such a way that the true age is contained randomly within this interval. A minimum interval of 0.0117 Ma (the shortest in the PBDB) was enforced, and ranges were checked not to cross the present. Time ranges were translated into uniform priors for the age of fossil terminals, an approach that has been found to improve tip-dated inference (Barido-Sottani et al. 2020). The age of extant terminals was fixed to the present.

The age of the root of the tree was treated as an unknown parameter, and priors for it were designed in a way that imitates the uncertainties of empirical phylogenetics. As routinely done, the earliest known fossil was used as a minimum age constraint. Note that this taxon is unlikely to be the true oldest member of the clade, as an average of 82% of extinct taxa were treated as unfossilized/undiscovered. If said taxon was not included in the final matrix (which only incorporated between 5 and 75 of the approximately 150 known fossil taxa, see above), then uncertainty in its age was generated as explained above and accounted for when establishing the minimum value for the root age prior. This parameter was modelled using both uniform and offset exponential distributions, allowing us to assess the effects of this decision. To find a justifiable width for these distributions, we scraped the online Fossil Calibration Database (https://fossilcalibrations.org/; Ksepka et al. 2015) using package *rvest* (Wickham 2021). We obtained 202 node calibrations with both minimum and maximum bounds, which showed an average difference of 107.7 Myr. When implementing a uniform root age prior, values were set to span 107.7 Myr prior to the first occurrence of the oldest known fossil lineage. When implementing an offset exponential distribution, the mean value was set so that 95% prior probability was contained in the 107.7 Myr preceding the minimum bound, thus establishing a less restrictive soft maximum (Yang & Rannala 2006, Warnock et al. 2011). These alternative root age priors are exemplified in Fig. S4.

Diffuse priors were used for most parameters of the birth-death model (exponential with a mean value of 10 for the speciation probability, default flat beta distributions for both extinction and fossilization probabilities), while the probability of sampling extant taxa was set to the true value. Morphological evolution was modelled using an Mkv+Γ model (Lewis 2001), accommodating both among-character rate heterogeneity and the ascertainment bias introduced by sampling only parsimony-informative characters. An uncorrelated relaxed morphological clock (IGR) was implemented (Lepage et al. 2007), with a variance drawn from an exponential prior with a mean value of 10. A normal distribution (mean=0.001, standard deviation=0.01) was used for the base clock rate, and the species sampling strategy was set to ‘fossiltip’ to avoid the recovery of fossils as direct ancestors, given that TREvoSim outputs reflect only cladogenetic processes. Two independent runs of four chains were continued for either 50 million generations or until attaining an average standard deviation of split frequencies (ASDSF) of 0.005, which was taken to represent a thorough sampling of posterior topologies (Mongiardino Koch et al. 2021). Trees were sampled every 1,000 generations, and a burn-in fraction of 0.5 was applied.

In order to assess the effect on node ages that is introduced by the models employed, we ran a subset of 350 analyses (50 from each level of fossil sampling, randomly selected) in the absence of morphological information. To this effect, we fixed the entire topology to match that of the simulated tree, obtaining dates that are informed exclusively by the ages of fossil tips and the model priors (Warnock et al. 2015, Brown & Smith 2018).

### Data analyses

Posterior distributions of topologies were imported into R and analyzed using packages already mentioned, as well as functions from *castor* (Louca & Doebeli 2017), *ggplot2* (Wickham 2016), *phangorn* (Schliep 2011), *readr* (Wickham & Hester 2020) and *stringr* (Wickham 2019). Two aspects of these topologies were explored through comparison with the true (i.e., simulated) tree: the placement of fossil taxa relative to the extant subtree, henceforth topological accuracy; and the inferred ages of the nodes in the extant subtree, henceforth temporal accuracy. In order to ensure uncertainty was being meaningfully captured, analyses with posterior distributions containing fewer than 100 trees were excluded. This occurred in a small number of replicates that included the fewest number of fossil terminals, as the extensive topological constraints meant low ASDSF values were reached rapidly.

Topological accuracy was explored by iterating through every fossil, and pruning true and posterior trees to just the extant taxa plus the focal fossil terminal. True and inferred fossil positions were compared in a random sample of 100 posterior trees using two approaches. First, we counted the fraction of posterior trees in which the fossil was placed correctly. Second, we calculated the average distance between true and inferred placements, estimated using the number of nearest-neighbour interchange (NNI) moves between the topologies. Note that some types of topological errors are not captured by these estimates, such as those relating to the relationships among entirely extinct clades. This was purposefully done in order to focus on metrics of topological accuracy that only capture the extent to which fossil tips are misinforming node ages in the extant subtree. The relative age of fossil terminals (i.e., their true age divided by the root age) and their proportion of missing data were gathered as potential predictors. To characterise the overall topological accuracy of each analysis, values for all fossil terminals were averaged.

Temporal accuracy was estimated in random samples of 1000 posterior trees pruned to just the extant taxa. For every node, we recorded the difference between the true age and the median posterior estimate, as well as whether the true age was contained within the 95% highest-posterior density (HPD) interval (known as the 95% HPD coverage). Relative errors were further obtained by dividing age differences by the true age value. Depending on the analysis, errors were expressed as signed deviations, or transformed into absolute values. To characterise the overall temporal accuracy of each analysis, values for all nodes were averaged.

The relationships between different estimates of topological accuracy and temporal accuracy were explored using non-parametric (Spearman’s rank) correlation coefficients. Generalised linear models incorporating the number of fossil terminals as an additional discrete predictor were also fit. Before these analyses, multivariate outliers whose squared Mahalanobis distance exceeded the 95% quantile value of a chi-square distribution were excluded. P-values were corrected for multiple comparisons using the Benjamini & Hochberg (1995) approach.

## Results

The accuracy with which fossils were placed in the extant scaffold was affected by both their age and proportion of missing data (Fig. 1A). The position of fossils was correctly inferred more frequently for fossils lying approximately half-way between the root of the tree and the present, but decreased for fossils that were younger and older. Missing data also negatively impacted the probability of correct fossil placement. However, accuracy remained high overall: only highly fragmentary fossils occupying both temporal edges of the tree (e.g., those within the earliest and latest decile) were incorrectly placed relative to their extant counterparts more often than not.

**Figure 1:**
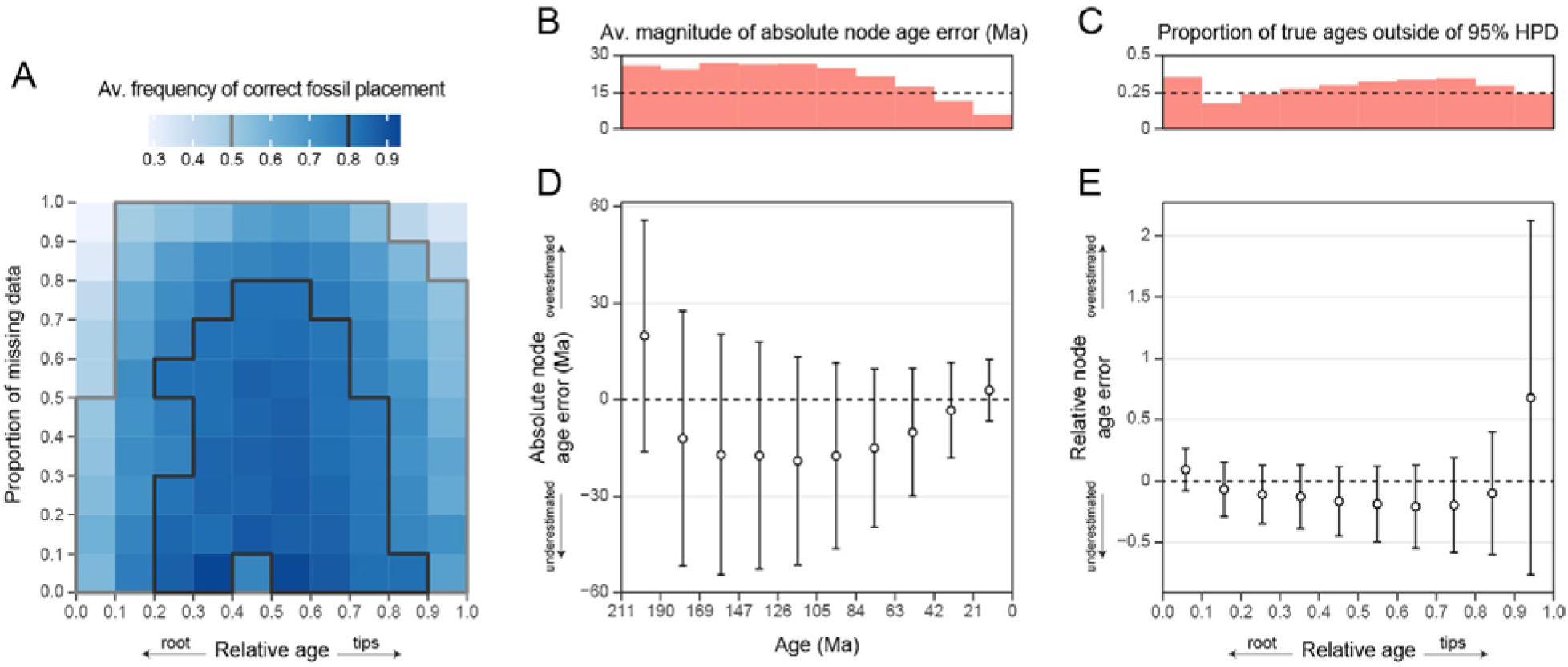
Determinants of accuracy in fossil placement and node age estimates. **A.** The frequency of correct fossil placement for various relative ages and proportions of missing data. Fossils with small amounts of missing data, and occurring further from the root and tips of the tree, are placed with higher levels of accuracy. **B.** Average magnitude of the absolute error in age estimates for nodes at different depths. Error increases with node depth before plateauing halfway between the root and tips of the tree. **C.** Proportions of nodes with true ages outside the 95% HPD. Values oscillate around 25%, with the highest fraction seen close to the root of the tree. **D.** Average error in absolute node age estimates. Values correspond to those shown in **B**, but averaged without removing the sign of the deviation, showing systematic biases in the direction of the estimation error. Circles denote the mean difference between true ages and median posterior estimates, error bars show the average width of the 95% HPD. Ages for the oldest and youngest nodes (i.e., those towards the root and tips) are generally overestimated, whilst those towards the middle of the tree tend to be underestimated. **E.** Relative error in age estimates for nodes at different depths. Values were computed as in **D**, but error was standardised by dividing by the true ages. Values correspond to analyses under an offset exponential prior for the root age; results under a uniform prior can be found in Fig. S8.

On average, error in the estimation of divergence times increased with node age up until half-way between the root and tips of the tree and remained constant thereafter, attaining absolute errors between 24.3 and 27 Myr (Fig. 1B). This corresponds to between 11.5-12.8% of total tree depth. When the sign of these deviations is considered, a convex relationship with age emerges, showing a complex interplay between accuracy and bias (Fig. 1D-E; a somewhat similar pattern can also be seen for analyses run under the joint prior, see Fig. S5). On average, divergence times for nodes lying close to both the root and tips of the tree were systematically overestimated, while all others were underestimated. Absolute error was highest for the deepest nodes, whose inferred ages exceeded true ages by an average of 19.8 Myr (Figs. 1D and S6); relative error was highest for the youngest nodes, whose inferred ages showed an excess of 68% of the true value (despite an average overestimation of only 2.9 Myr; Fig. 1E). While these two regions of the tree also showed the highest levels of error in fossil placement (Fig. 1A), we note that temporal windows with high topological accuracy (i.e., those towards the centre of the time spanned by the phylogeny) still had relatively large absolute levels of temporal error. Despite this, overall accuracy was also relatively high, with 95% HPD intervals including true ages 65.2-83.5% of the time, with the lowest values seen among the nodes closest to the root (Fig. 1C). While Figure 1 shows only results using offset exponential distributions for the root age prior, nearly identical results were obtained when implementing uniform root age priors (Fig. S7). The only noticeable difference was a slightly higher proportion of true ages outside the 95% HPD (i.e., lower coverage) for nodes closer to the root of the tree (Fig. S8).

The overall temporal and topological accuracy of analyses (obtained by averaging the data for all node ages and fossil placements) increased with the number of fossils sampled (or, conversely, their overall temporal and topological error decreased; see Fig. S9). While the increase in temporal accuracy likely reflects the positive influence of a larger number of fossil tips informing node ages, the increase in topological accuracy is probably related to the reduced number of branches in the extant subtree to which fossils can attach, as well as their increased length (Fig. S10), that result from a sparser sampling of extant taxa.

If not accounted for, the effect of the level of fossil sampling on both topological and temporal accuracies confounds a direct exploration of the relationship between the two variables. We took two approaches to decouple these effects. First, the data were subdivided by the different levels of fossil sampling, and correlations between temporal and topological accuracy were explored within each subset. Overall, correlations were weak (absolute p values ≤ 0.29), inconsistent in sign (with 5 instances of positive and 9 of negative correlations), and non-significant in 12 out of 14 subsets (Fig. 2). The only two examples of significant correlations between these variables involved negative correlation coefficients, contrary to our expectations (Fig. 2). A second approach involved including both the level of fossil sampling and the topological accuracy as predictors of temporal accuracy in a generalised linear model. Once again, while topological accuracy was recovered as a significant predictor (P = 0.003 and 0.007, depending on the proxies of accuracy employed), its effects were relatively small and negative in sign (estimates = −0.08 and −0.06, respectively). The small impact of topological accuracy on temporal accuracy is further reflected by the fact that including this predictor alongside the level of fossil sampling in the linear model only increased the adjusted R^2^ values by 0.004 to 0.009 (relative to R^2^ values between 0.204 and 0.291 when using the level of fossil sampling as sole predictor). Once again, these results are not affected by the root age prior implemented (see Fig. S11 for results under a uniform prior).

**Figure 2:**
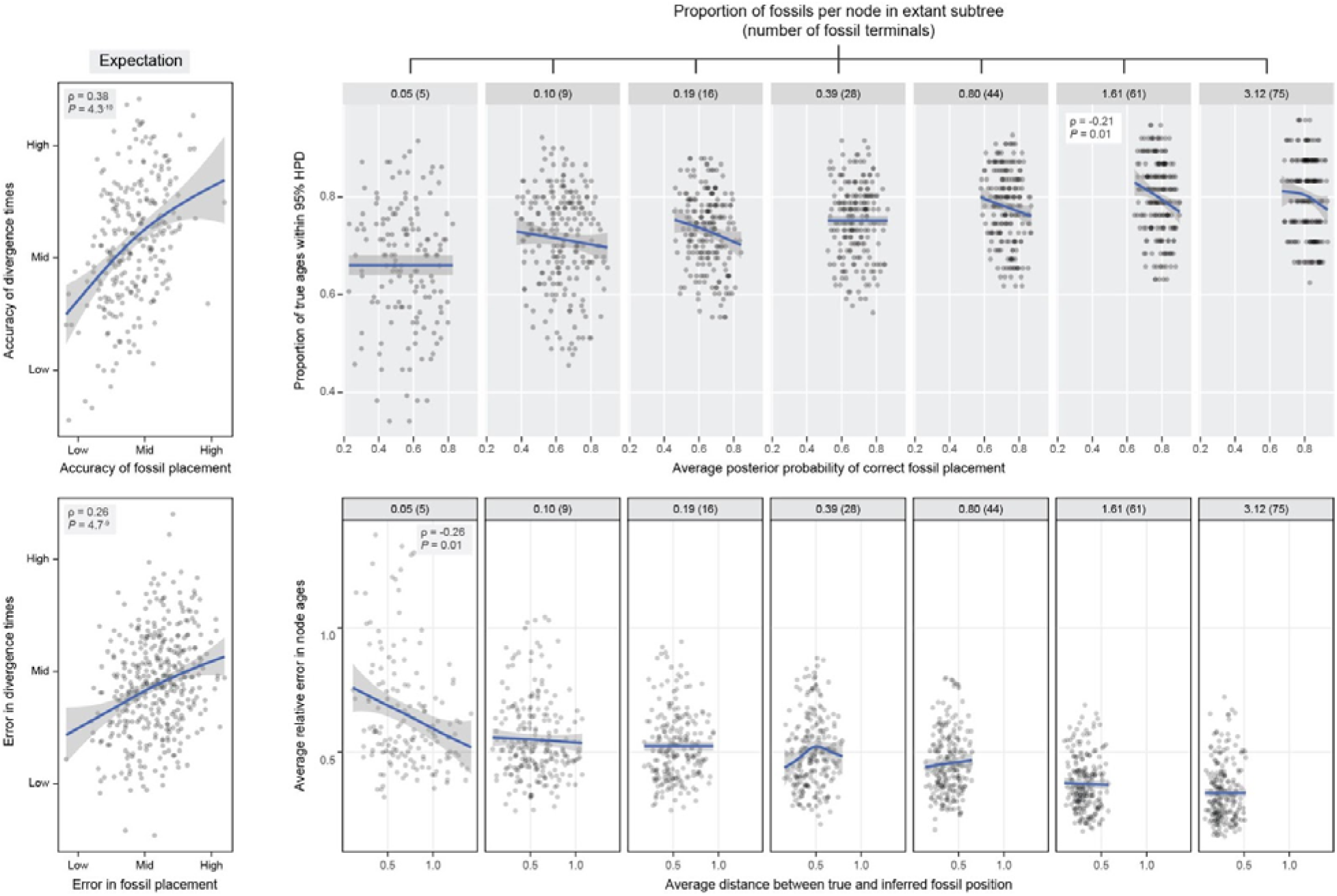
Higher topological accuracy does not translate into higher temporal accuracy. Top and bottom graphs show alternative proxies to estimate the success in inferring overall fossil placement and divergence times. Within each plot, a positive relationship is expected under the hypothesis that accurate fossil placement improves divergence time estimates (shown left). No such relationship is obtained, with topological accuracy being either uncorrelated or negatively correlated with temporal accuracy (ρ and P values correspond to Spearman’s rank correlations). Trend lines were fitted using general additive models, and are shown for visualisation purposes only. Values correspond to analyses under an offset exponential prior for the root age; results under a uniform prior can be found in Fig. S11.

## Discussion

Phylogenetic trees with optimal levels of fossil sampling are key to reconstructing evolutionary history (Quantal & Marshall 2010, Slater & Harmon 2013, Rothwell et al. 2018), and have illuminated the diversification of clades in ways unattainable by studies focusing only on living representatives (Slater 2013, Finarelli & Goswami 2013, Garwood et al. 2014, Betancur-R et al. 2015, Mitchell 2015, Arcila & Tyler 2017, Vinther et al. 2017, Norrell et al. 2020, Lloyd & Slater 2021, Mongiardino Koch & Thompson 2021, Wisniewski et al. 2022). The macroevolutionary potential of the fossil record is best unlocked through the use of tip-dated methods of inference that place fossil and living taxa in a common time-calibrated phylogenetic framework, built using mechanistic models of diversification and sampling, and informed by all available sources of information (Ronquist et al. 2012a, Gavryushkina et al. 2014, Zhang et al. 2016, Heath et al. 2017, Warnock & Wright 2021). One of the major benefits of this approach is that it facilitates the inclusion and extensive use of paleontological data (O’Reilly et al. 2015, Donoghue & Zhang 2016, O’Reilly & Donoghue 2020). Nonetheless, for these benefits to be realised, morphology must retain some degree of temporal signal (i.e., ‘morphological clocks’ need to exist, at least to some degree; Polly 2001, Beck & Lee 2014, Lee et al. 2014, King et al. 2017, Parins-Fukuchi & Brown 2017, Barba-Montoya et al. 2021), and morphological data need to allow inference of the phylogenetic position of fossil terminals (Sansom et al. 2010, Sansom & Wills 2013, Guillerme & Cooper 2016, Luo et al. 2020, Mongiardino Koch et al. 2021).

Our results show that morphological datasets of moderate sizes (300 characters) and with realistic levels of missing data generally succeed in inferring both the position of fossil terminals and the divergence times of extant lineages, at least when combined with topological constraints for living terminals. The correct position of fossils relative to fixed extant scaffolds is generally attained with high confidence, although this is impaired by high levels of missing data, and is particularly difficult for both very ancient and very recent fossil taxa (Fig. 1A). These patterns are also seen in empirical morphological datasets, which generally target divergent taxa to avoid issues of poor resolution, and often exhibit poor accuracy for deep splits (Scotland et al. 2003, Ronquist et al. 2012a, Pyron 2015, Halanych 2016, Ronquist et al. 2016, Zhang et al. 2016). Similarly, despite an increase in the error of inferred node ages towards the root of the tree (Fig. 1B), true ages are generally included within the 95% HPD intervals that are commonly used to summarise posterior distributions of trees (Fig. 1C), revindicating the value of morphological clocks for dating phylogenies.

Despite the overall high levels of accuracy, we also uncover systematic biases in the inference of divergence times (Figs. 1D-E and S7D-E). These manifest in the overestimation of node ages close to both the root and the tips of the tree, while the ages of all remaining nodes tend to be underestimated. Excessively old ages for deep nodes were also a commonly reported artefact of early tip-dating analyses (e.g., Ronquist et al. 2012a, Slater 2013, Arcila et al. 2015). A number of drivers were proposed for this ‘deep root attraction’ phenomenon, such as conflicts between morphological and molecular datasets, the use of inappropriate uniform tree priors, and a failure to accommodate for diversified sampling, or to account for morphological integration (Arcila et al. 2015, Ronquist et al. 2016, Zhang et al. 2016), but none of these apply to our simulations. Recently, Luo et al. (2020) suggested that instances of overestimation of deep node ages might be linked to fossil misplacements. Our analysis also finds node ages to be overestimated in regions of the tree where fossil placement is most inaccurate. Even so, regions of the tree where fossils tend to be placed with high accuracy still show relatively large absolute errors as well as biased estimates, although in this case the tendency is towards recovering systematically younger dates. Therefore, these effects seem unlikely to be driven by a failure to correctly place fossils, but rather by a mismatch between the constant-rate FBD process implemented and the overall shape of our phylogenetic trees (Figs. S5-S6), possibly coupled with the limited capability of small amounts of morphological information to overrule model priors. In fact, previous studies that simulated topologies under birth-death processes found little to no overestimation of the age of deep nodes (Zhang et al. 2016, Luo et al. 2020), which further highlights the importance of simulation approaches that incorporate some degree of model misspecification, as is expected of empirical datasets (O’Reilly et al. 2016, Ronquist et al. 2016, Goloboff et al. 2018, Mongiardino Koch et al. 2021). If these biases do arise from model misspecification, they might not affect tip dating in general, and only impact clades that have undergone ancient and rapid radiations (see also Budd & Mann 2020). For such clades, it remains possible that the addition of molecular data would succeed in overcoming these biases, although divergence times derived from molecular data can also be highly sensitive to model priors (see e.g., Warnock et al. 2015, Brown & Smith 2018, Mongiardino Koch et al. 2022, Sauquet et al. 2022). Alternatively, skyline FBD models that accommodate changes in rate parameters through time should also be considered when there is sufficient *a priori* information that diversification or sampling rates have been variable (Zhang et al. 2016, Simões et al. 2020, May et al. 2021, Wright et al. 2021, Barido-Sottani et al. 2022). Still, studies inferring tip-dated trees under a morphological clock should be aware that node ages might be strongly determined by the choice of tree prior, and that any given FBD model can still impose biases if it fails to capture relevant aspects of the history of diversification (Ronquist et al. 2016, Matschiner 2019).

While previous studies had cautioned that correct fossil placement was likely necessary to infer accurate divergence times (O’Reilly et al. 2015, Donoghue & Zhang 2016, Luo et al. 2020), we find no evidence for this hypothesised effect. Although extremely inaccurate fossil placements have been shown to distort divergence time analyses (Lee 2009, Near et al. 2005, Barido-Sottani et al. 2022), tip-dating using medium-sized morphological datasets, and implementing extant scaffolds, is sufficiently robust to result in broadly accurate timetrees. Furthermore, divergence times significantly improve with increased fossil sampling, but are not strongly impacted by the success with which fossil tips are placed relative to extant taxa (Figs. 2 and S11). This result is encouraging. It proves that a thorough sampling of fossils is beneficial for tip dating, even when many of these taxa prove unstable, or even resolve in slightly incorrect positions. Palaeontologists should be aware of the general difficulties faced when attempting to infer the relationships of ancient, young, and highly incomplete fossil terminals, but their *a priori* exclusion from tip-dated inference under morphological clocks is not justified. When coupled with molecular scaffolds for extant lineages, tip dating benefits from the advantages of co-estimating tree topology and divergence times, without suffering from its potential drawbacks.

## Acknowledgements

Analyses were run using the HTCondor service provided by the IT Services Research Infrastructure team at the University of Manchester, which runs the HTCondor software developed by the CHTC Team at UW-Madison, Wisconsin. The HTCondor service also utilises the AWS Marketplace Community AMI HTCondor image used for “bursting into the (AWS) cloud”. The manuscript benefited from discussions with Dahiana Arcila.

## Supplementary Figures

**Figure S1:**
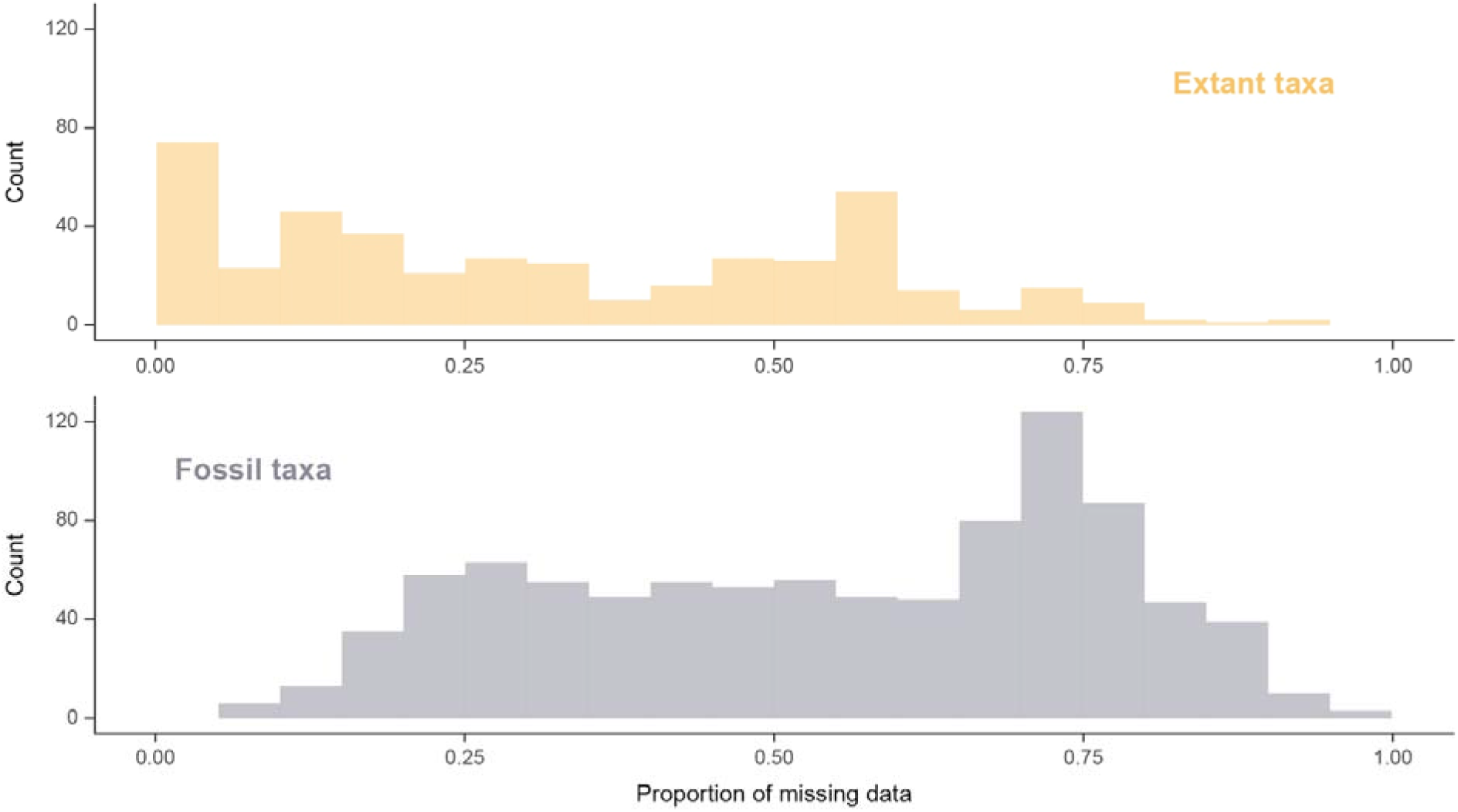
Empirical distributions of the proportion of missing data across extant and fossil taxa. Data was gathered from the same 12 empirical datasets employed to validate other aspects of the evolutionary simulations. See matrix-level distributions of missing data in Fig. S2.

**Figure S2:**
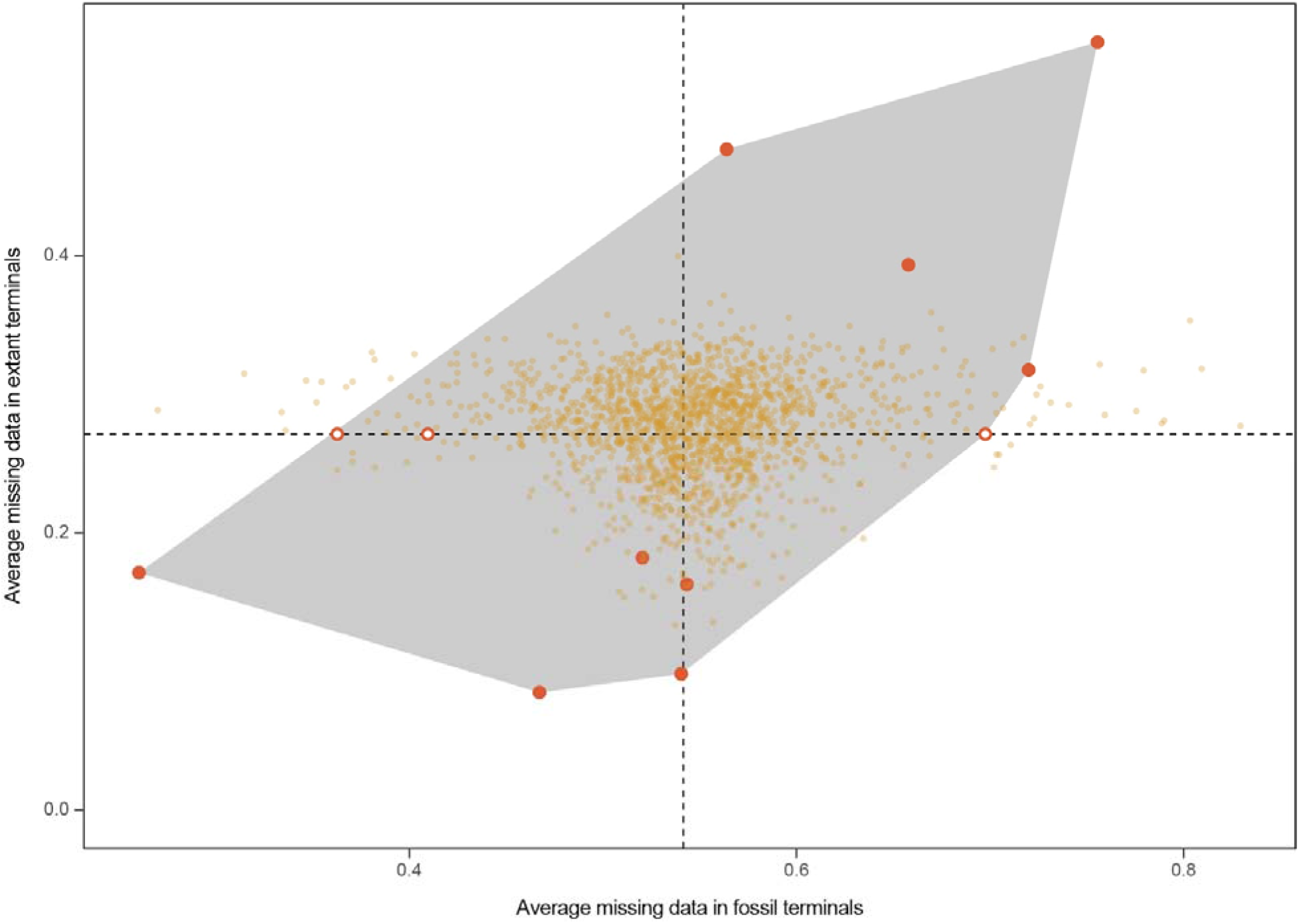
Average amounts of missing data across empirical (red) and simulated (yellow) datasets. Axes correspond to the mean fraction of missing data per terminal, for both fossil and extant terminals (see taxon level distributions of missing data in Fig. S1). Dotted lines mark the location of the centroid for empirical datasets. The three empirical datasets shown with empty circles contained exclusively fossil terminals, and their location on the y-axis is therefore fixed to the mean of the remaining datasets. Over 98% of simulated datasets fall within the convex hull determined by empirical datasets, with only a handful of matrices showing excessively low/high amounts of missing data among fossil terminals.

**Figure S3:**
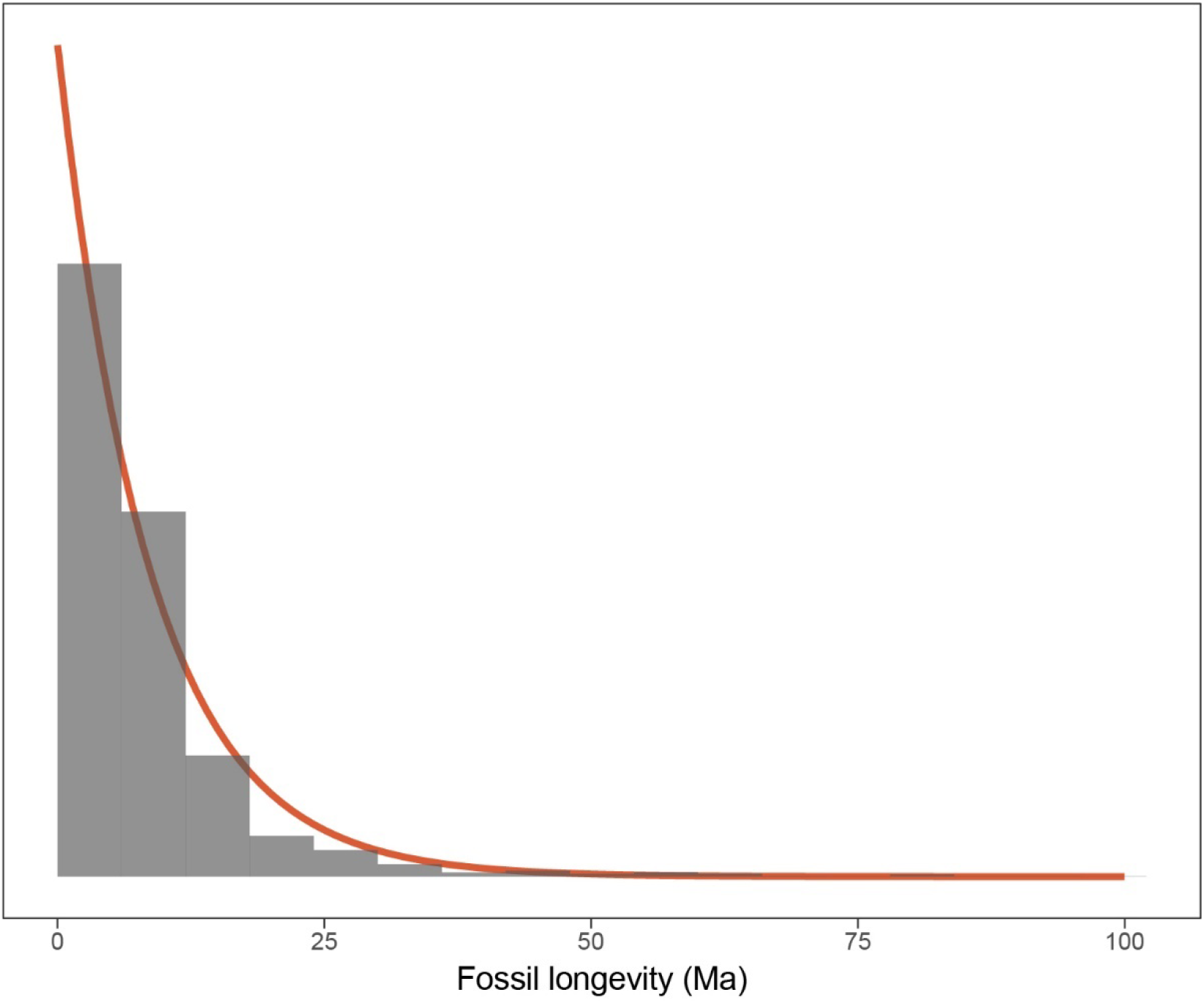
Empirical (grey bars) and fitted (red line) distributions of fossil longevities in millions of years (Ma). Empirical data corresponds to the entire Paleobiology Database (PBDB, https://paleobiodb.org/), truncated to a maximum longevity of 100 Ma. Fitted line corresponds to an exponential distribution, with an offset of 0.0117 Ma (minimum value observed in the data). Mean longevity of both distributions is 8.67 Ma.

**Figure S4:**
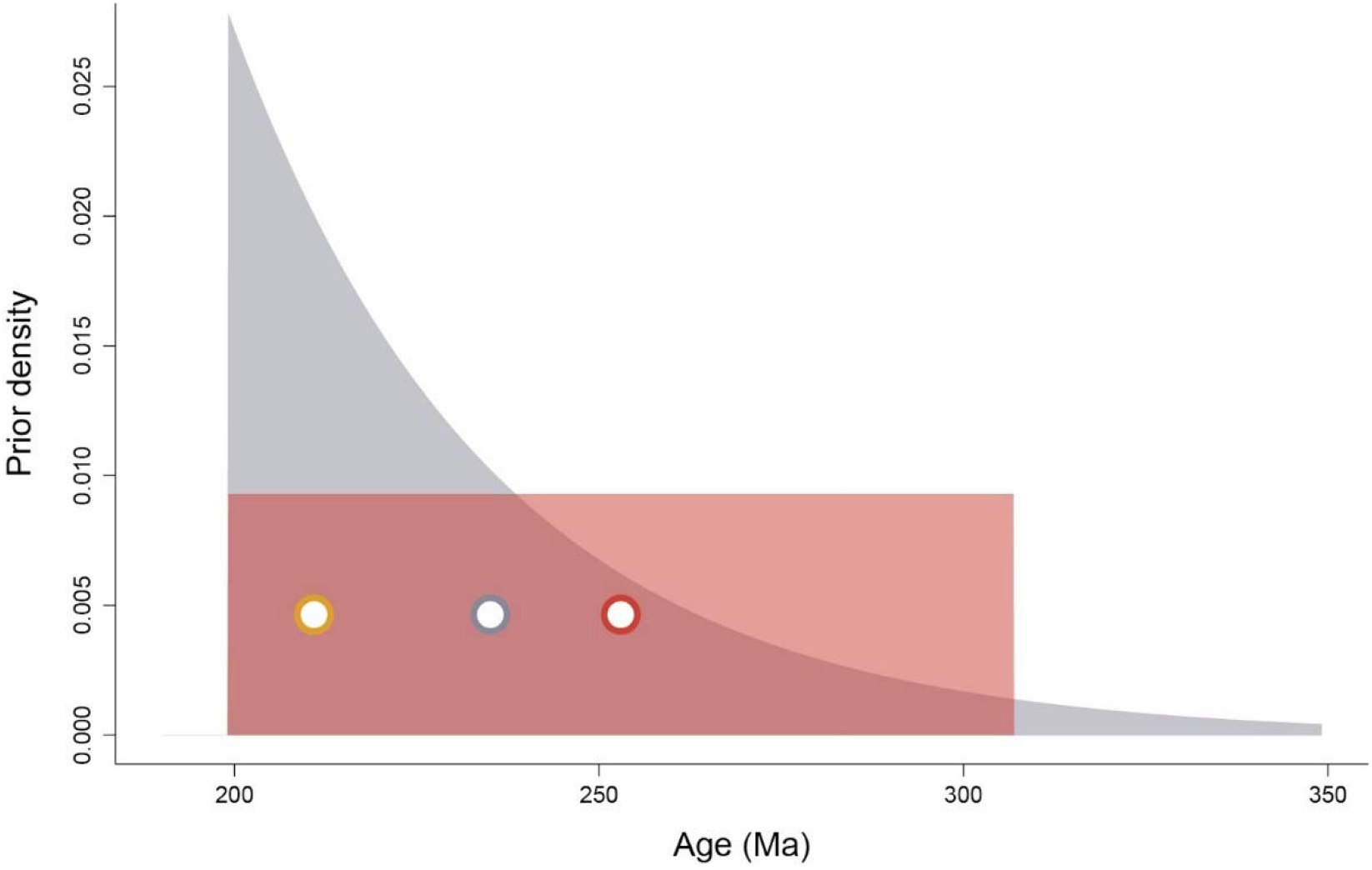
Alternative prior probabilities used to constrain the root age of the FBD analyses, exemplified with a dataset whose earliest fossil representative had an older bound at 199.13 Ma. Uniform prior (red) were designed to span a time frame extending 107.7 Myr into the past, while exponential priors (grey) contained 95% prior density within that same interval, but had a soft maximum bound that allowed for much older ages. Nonetheless, the mean values of exponential priors (grey circle) were always closer to the true age (yellow circle) than those of uniform priors (red circle)

**Figure S5:**
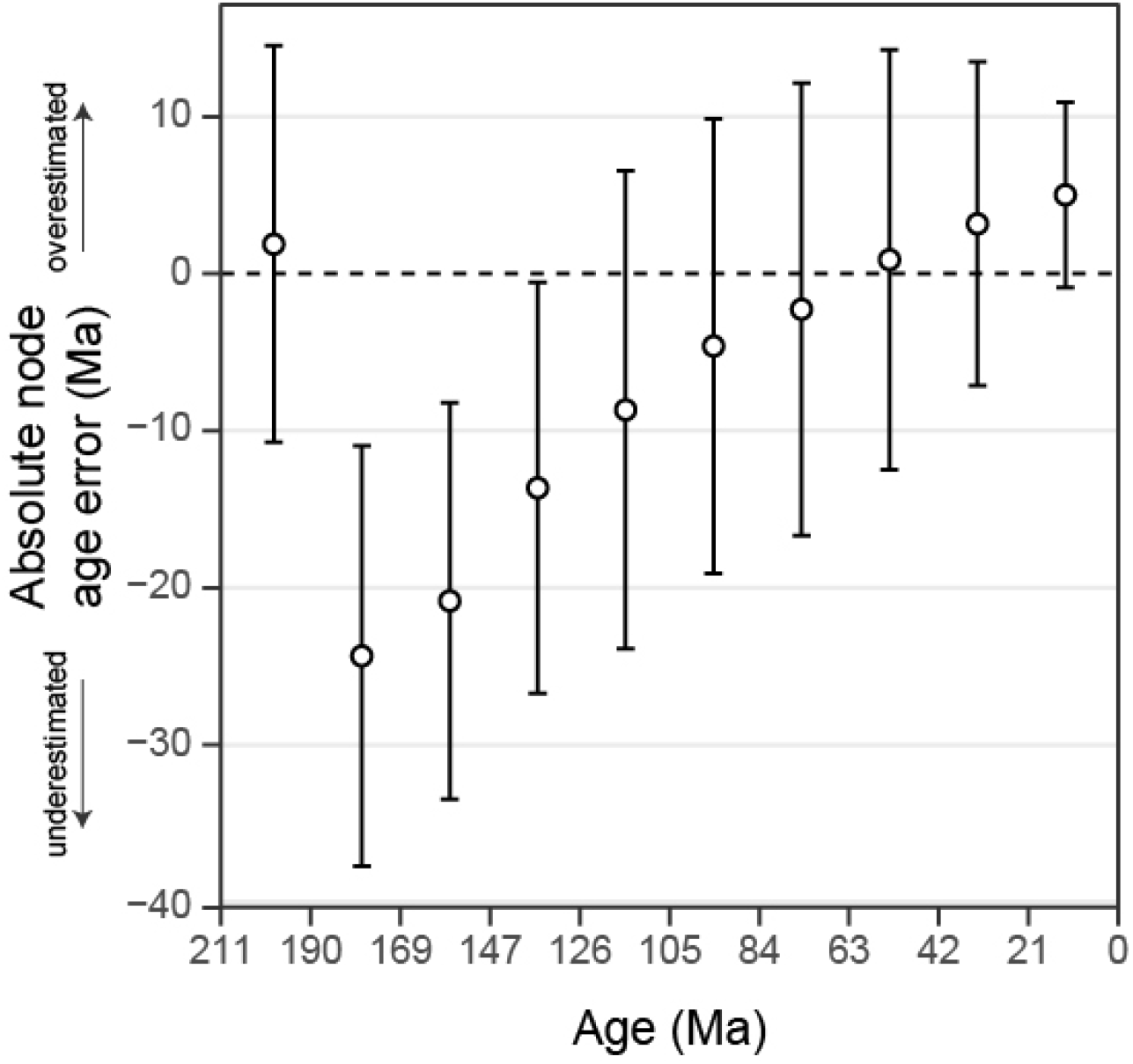
Average error in absolute node age estimates obtained under the joint prior. Similar patterns of systematic bias to those obtained in the full analyses (shown in Figs. 2D and S7D) can be observed, including a convex relationship between temporal bias and node depth, and a tendency to overestimate the age of young and old nodes, and underestimate those of all others. However, note that the absolute node age error of nodes closest to the root of the tree is much smaller than in the analyses with morphological data.

**Figure S6:**
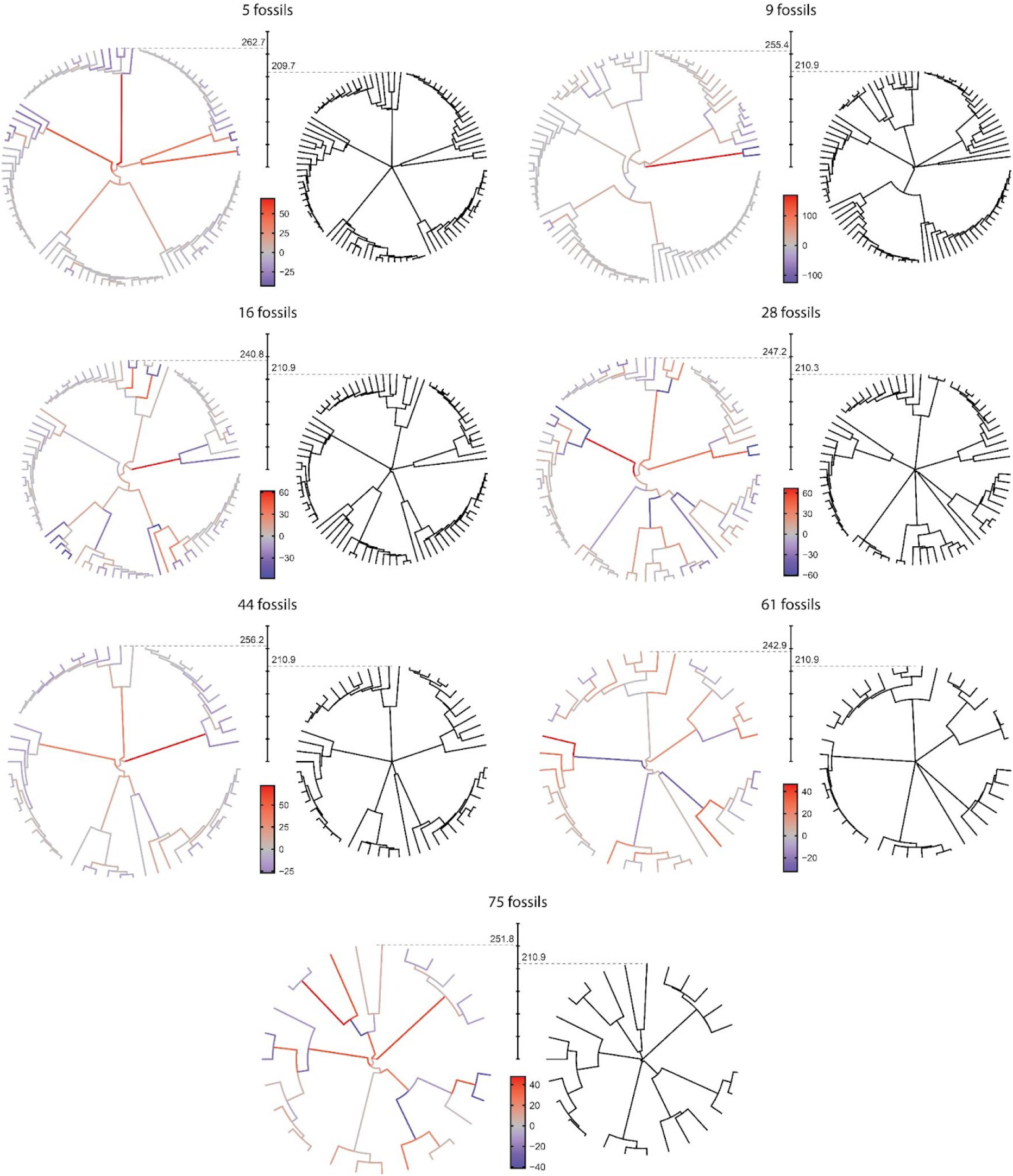
Examples of mismatches between inferred (left) and true (right) extant subtrees. One simulation per level of fossil sampling was selected at random, and a consensus extant subtree was built using median ages among posterior topologies. Branches are colour coded based on the level of over/underestimation of their lengths. The common temporal axis displays time since the origin, and has tick marks separated by 50 Ma. Excessively old ages for deep nodes are mostly obtained through the stretching of long branches connecting the clades originating from the deep ancient radiation, a pattern common to all levels of fossil sampling.

**Figure S7:**
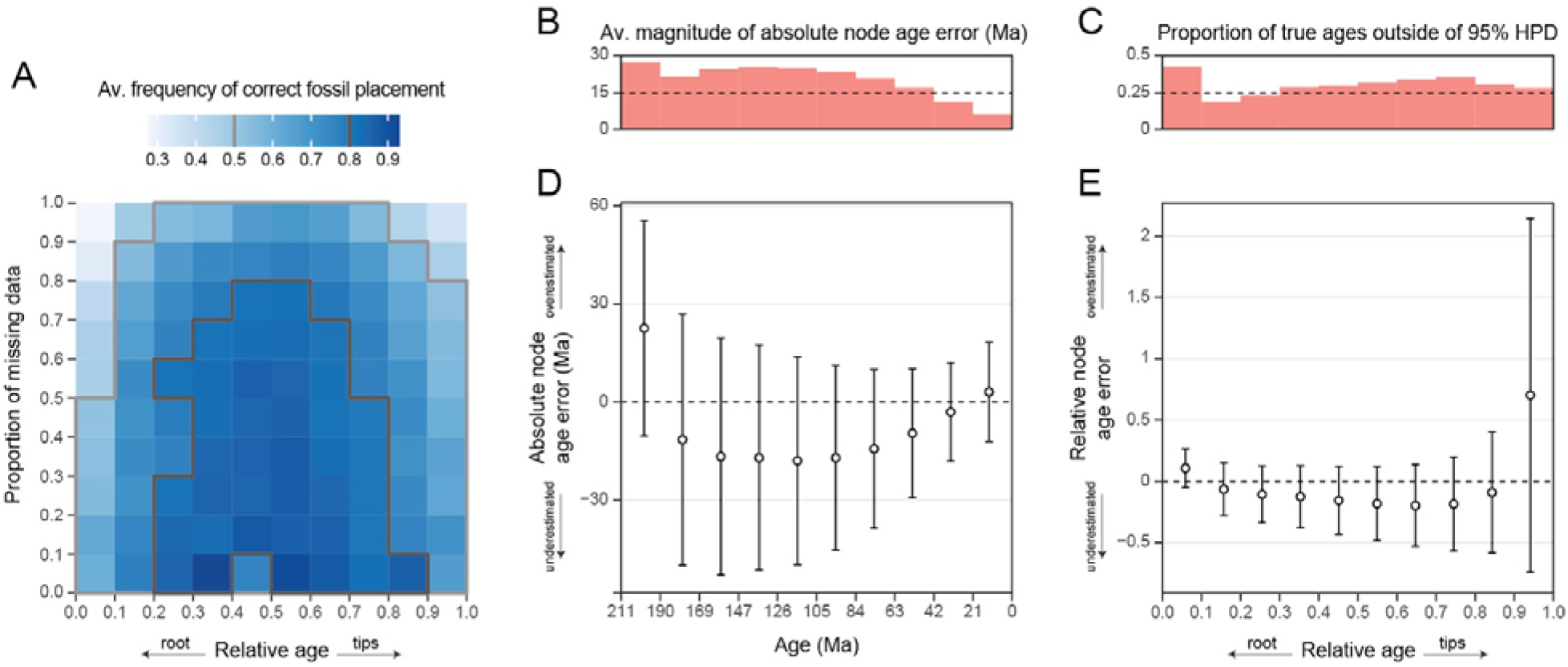
Determinants of accuracy in fossil placement and node age estimates. The variables depicted are equivalent to those of Fig. 1, but results correspond to analysis under a uniform root age prior. **A.** The frequency of correct fossil placement for various relative ages and proportions of missing data. **B.** Average magnitude of the absolute error in age estimates for nodes at different depths. **C.** Proportions of nodes with true ages outside the 95% HPD. **D.** Average error in absolute node age estimates. Values correspond to those shown in **B**, but averaged without removing the sign of the deviation, showing systematic biases in the direction of the estimation error. Circles denote the mean difference between true ages and median posterior estimates, error bars show the average width of the 95% HPD. **E.** Relative error in age estimates for nodes at different depths. Values were computed as in **D**, but error was standardised by dividing by the true ages.

**Figure S8:**
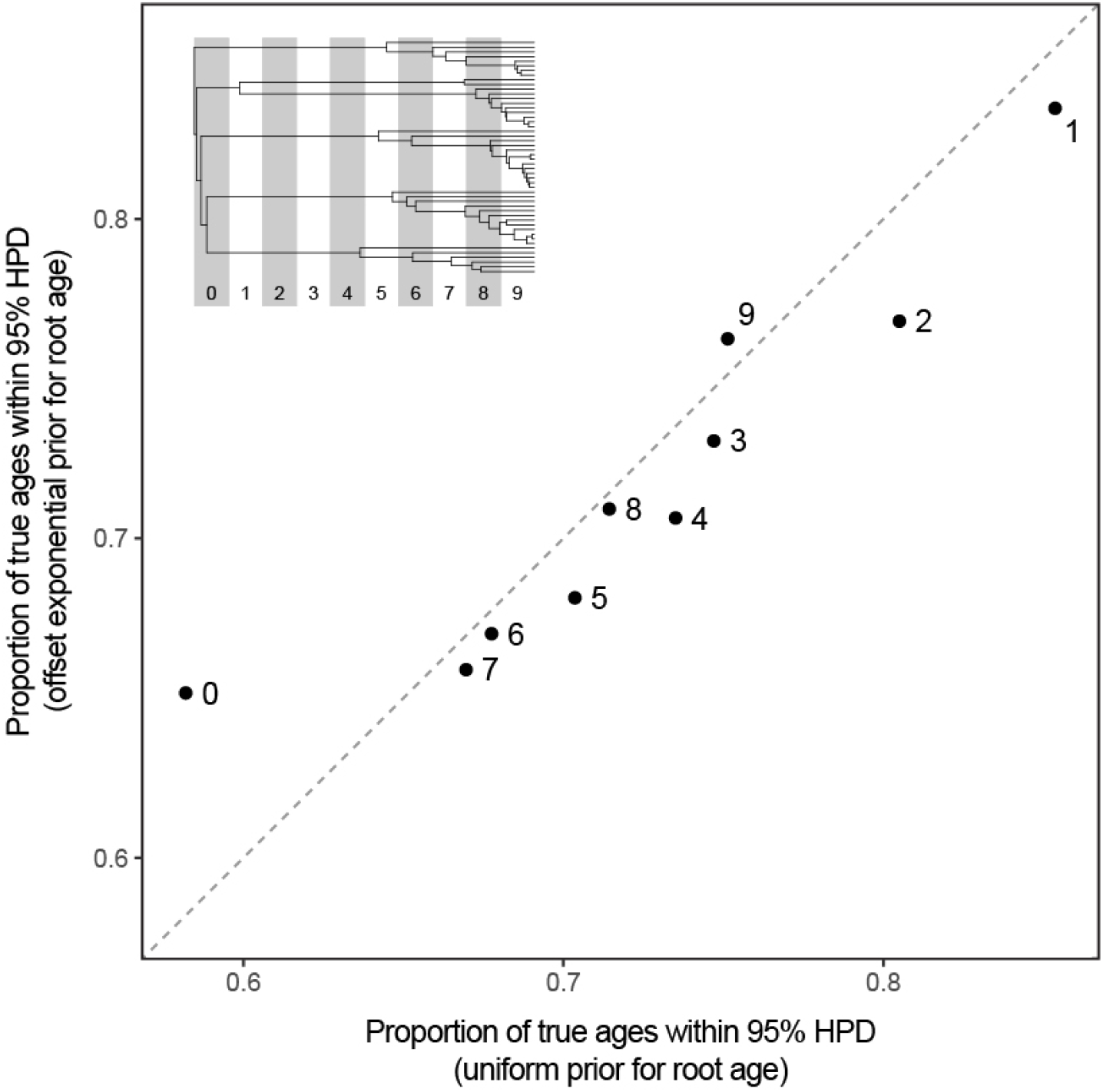
Comparison of node age coverage (i.e., fraction of true ages contained within 95% HPD inference intervals) for analyses run under uniform and offset exponential priors for the age of the root. For each analysis, coverage was estimated after binning true node ages into 10 intervals spanning the distance between root and tips (labelled as 0 and 9, respectively; see inset), and values averaged across all analyses. The dashed line shows equal coverage across both conditions. The largest deviation occurs for the oldest nodes, for which analyses under a uniform root age prior show lower coverage.

**Figure S9:**
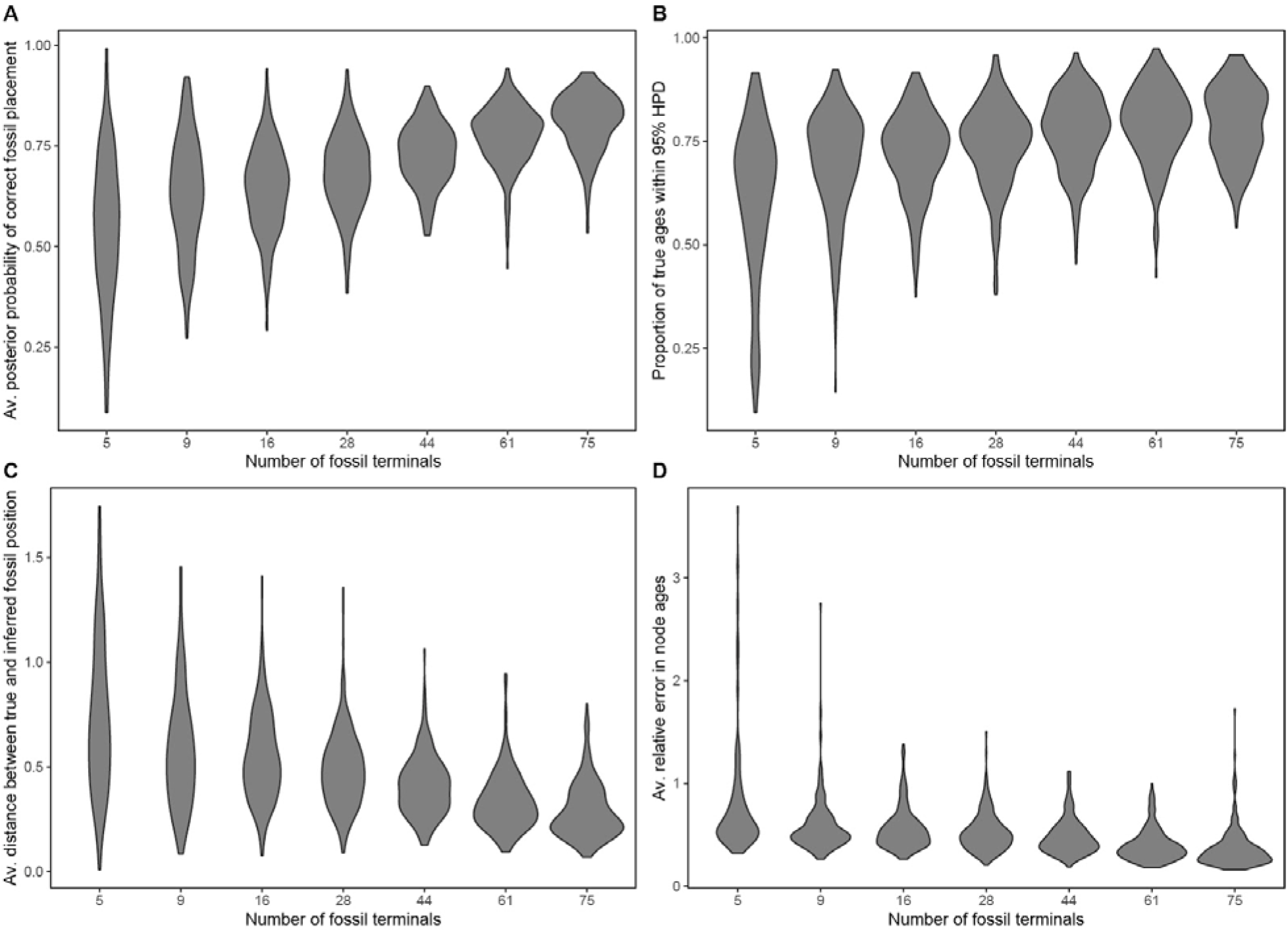
Increased fossil sampling results in higher topological accuracy (i.e., increased probability of inferring the correct position of fossil terminals [**A**], decreased distance between true and inferred positions [**B**]; see also Fig. S10) and higher temporal accuracy (increased proportion of true node ages contained within 95% HPDs [**B**], decreased error in node age estimates [**D**]). Results correspond to offset exponential distributions to model the root age prior, but results are identical under uniform priors.

**Figure S10:**
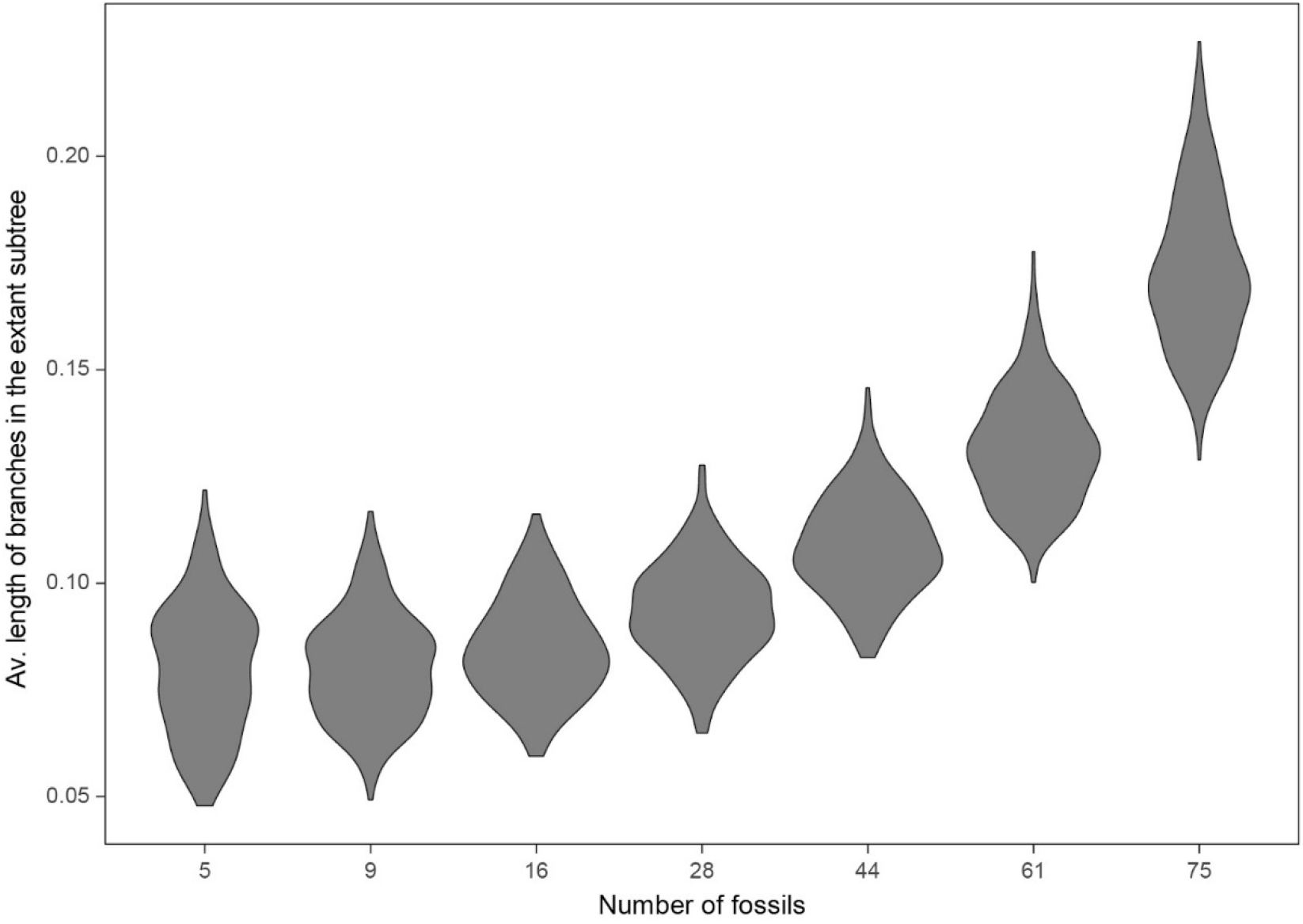
The average length of branches in the extant subtree increases as fossils represent a larger fraction of sampled terminals. This, plus the overall fewer number of branches in the extant subtree, likely drive the higher topological accuracy seen as more fossils are sampled (see Figs. 2 and S9). Branch lengths were standardised by the root-to-tip distance.

**Figure S11:**
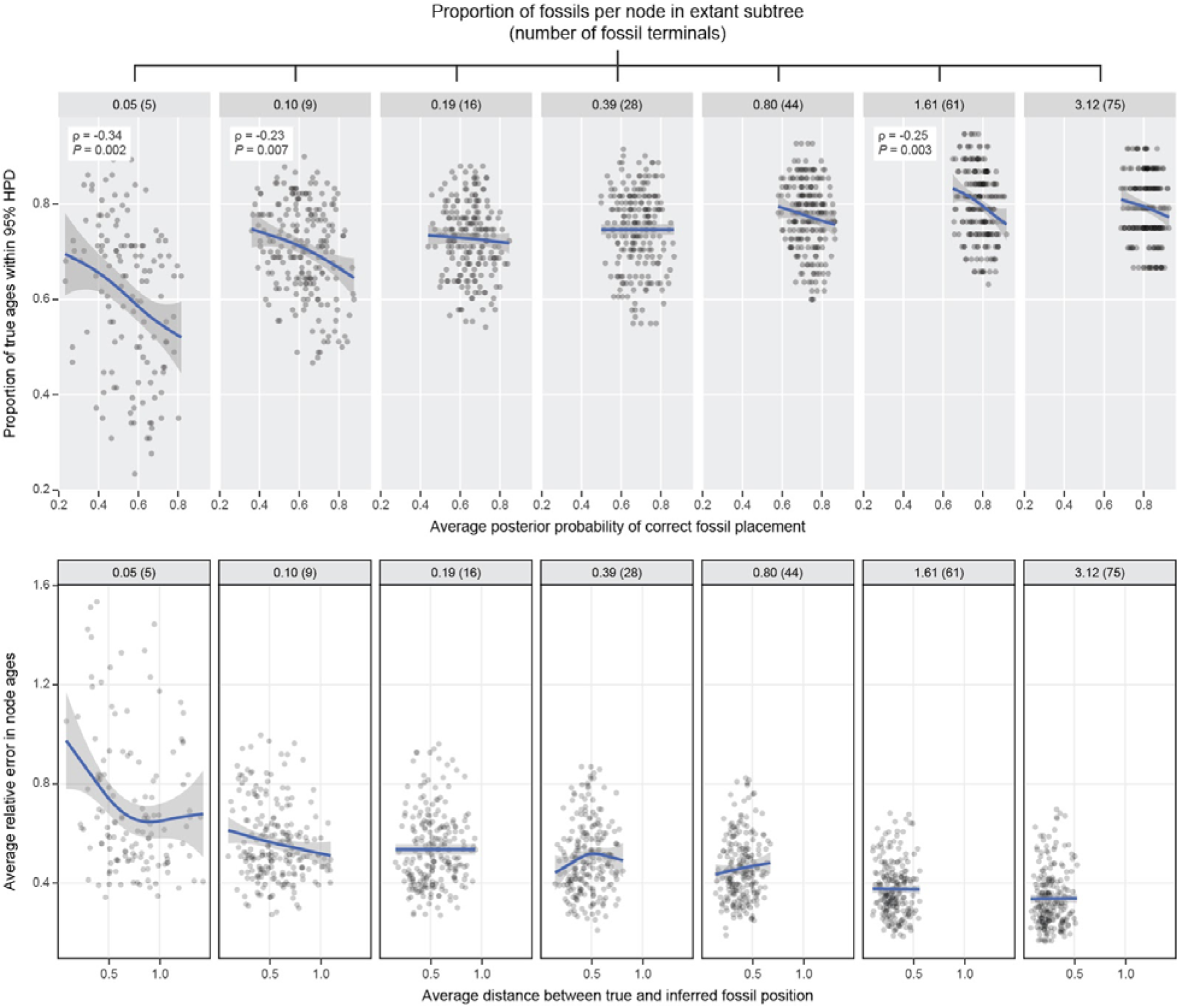
Higher topological accuracy does not translate into higher temporal accuracy.. The variables depicted are equivalent to those of Fig. 2, but results correspond to analysis under a uniform root age prior. Top and bottom graphs show alternative proxies to estimate the success in inferring overall fossil placement and divergence times. Relationships between these variables are never positive, as expected under the null hypothesis (see Fig. 2). Trend lines were fitted using general additive models, and are shown for visualisation purposes only; ρ and P values correspond to Spearman’s rank correlations. The addition of topological accuracy to the generalised linear models increased R^2^ values from 0.271-0.316 to 0.301-0.318, relative to models including only the level of fossil sampling as predictor.

